# Quantification of Antisense Oligonucleotides by Splint Ligation and Quantitative Polymerase Chain Reaction

**DOI:** 10.1101/2021.06.05.447195

**Authors:** Minwook Shin, Pranathi Meda Krishnamurthy, Jonathan K. Watts

## Abstract

Reliable detection and quantification of antisense oligonucleotides (ASOs) in experimental and clinical specimens is essential to understand the biological function of novel oligonucleotide-based therapeutics. In this study, we describe a method to detect and quantify ASOs in biological samples, whereby the ASO acts as a splint to direct the ligation of complementary probes and quantitative real-time PCR was used to monitor ligation products. Low levels of 2′-*O*-MOE gapmer ASO in serum, liver, kidney, lung, heart, muscle, and brain tissues can be detected over a 6-log linear range for detection using this method. This method allows quantification of various types of chemically modified ASOs, including PS linkage, 2′-OMe, 2′-*O*-MOE, locked nucleic acid (LNA), and siRNA. This method does not require probe modifications, and can be performed using standard laboratory equipment; making it a fast, sensitive, and reliable technique that can be widely applied. This detection method may find potential applications in detection of therapeutic oligonucleotides in biological samples.

## 1. Introduction

Antisense oligonucleotides (ASOs) consist of DNA, RNA, or chemically modified nucleotides that regulate target RNA molecules through Watson-Crick base pairing [1-3]. For potential therapeutic applications the physiological stability and pharmacokinetics of ASOs need to be optimized by introduction of chemical modifications to the phosphate linkage, ribose sugar, or nucleobase [4-6]. The phosphorothioate (PS) linkage involves the replacement of a non-bridging oxygen in the phosphodiester linkage with sulfur which improves nuclease resistance and facilitates cell uptake through increased protein binding [7,8]. Modification of the ribose sugar with 2′-*O*-methyl (2’-OMe) RNA [9], 2′-*O*-methoxyethyl (2′-*O*-MOE) RNA [10], or locked nucleic acid (LNA) [11] improves affinity towards the target RNA, and increases the stability of the ASO by protecting it from nuclease digestion. When fully modified ASOs tightly bind to target RNAs they function as steric blockers to regulate functions such as splicing [12,13]. Gapmers are ASOs which typically have a stretch of phosphorothioate-modified DNA in the central region to recruit RNase H, usually flanked by high affinity, nuclease-stable modified nucleotides [14]. All of these modifications increase the potential use of ASOs as therapeutic agent in the treatment of life-threatening diseases. Novel chemical modifications, improved knowledge of mechanisms of action, and refined clinical trial designs have all contributed to increasing momentum for the translation of ASOs into the clinic [15]. Indeed, more than 300 ASOs are now in various stages of late preclinical and clinical development [16].

Simple and reliable bioanalytical methods to detect and quantify ASOs are essential to understand their mechanism of action and to transition research from preclinical to clinical settings [17]. The ASO detection methods being used currently vary in the costs incurred for sample preparation and analysis, as well as their sensitivity. Liquid chromatography coupled with mass spectrometry (LC-MS) can discriminate ASOs from their metabolites, but require extensive sample processing, and the sensitivity of the method is relatively low, requiring ASO concentrations above 10 pM [18,19]. Hybridization-based enzyme linked immunosorbent (ELISA) assays have a higher sensitivity (100 fM) and can be performed with moderate throughput [20,21]; however, their linear range for detection is low, and the extensive sample processing required makes the approach time-consuming and expensive.

The quantitative real-time Polymerase Chain Reaction (qPCR) has the highest sensitivity and broadest detection range. However, because of the short length of ASOs, it is rather challenging to apply qPCR to their quantitation [22]. Increasing the lengths of qPCR templates can address this limitation. For example, stem-loop reverse transcription (RT)-qPCR works well for quantitation of ASOs with simple modifications, but performs poorly with ASOs with complex modifications, like fully modified 2′-*O*-MOE [23]. Therefore a chemical ligation qPCR method using chemical ligation between modified oligonucleotide probes, which have 3′-phosphorothioate and 5′-phenylsulfonyl groups, was developed [23]. However, this chemical ligation qPCR method requires oligonucleotide probes with modified 3′ and 5′ ends which are neither commercially available nor easy to produce in most laboratories. A similar splint-ligation approach mediated by *Chlorella* virus SplintR^®^ DNA ligase (New England Biolabs Inc., Ipswich, MA, USA) has previously been reported to be used for miRNA quantification [24,25]. The SplintR qPCR assay is a quick and cost-effective method that can be used in any laboratory. In this study, we show that the SplintR qPCR assay effectively detects and quantifies ASOs in biological samples.

## 2. Materials and Methods

### 2.1. Synthesis of ASOs

Short interfering RNAs were purchased from Integrated DNA Technologies (IDT, Coralville, IA, USA). ASOs were synthesized in-house using Applied Biosystems 394 DNA/RNA synthesizers with UnyLinker CpG supports (ChemGenes, Wilmington, MA, USA) and standard detritylation and capping reagents, as described previously [26]. Activation involved the use of 5-benzylmercaptotetrazole (0.25 M in acetonitrile, ChemGenes), and oxidation using iodine (0.05 M in a 9:1 mixture of pyridine:water, ChemGenes), as well as sulfurization with 3-((dimethylamino-methylidene)amino)-3H-1,2,4-dithiazole-3-thione (DDTT) (0.1 M, ChemGenes). DNA, 2′-OMe, 2′-*O*-MOE, and LNA phosphoramidites (ChemGenes) were dissolved in anhydrous acetonitrile to a concentration of 0.15 M. LNC-C phosphoramidite was dissolved in THF:acetonitrile (3:1). 2′-OMe-U phosphoramidite was dissolved in acetonitrile:DMF (8.5:1.5). Modified phosphoramidites were coupled for 10 min. The ASOs were deprotected and cleaved from the CpG support using concentrated NH_4_ OH for 16 h at 55°C. Oligonucleotides were desalted by ultrafiltration on Amicon Ultra Centrifugal Filters (Millipore, St. Louis, MO, USA). All ASOs were characterized by electrospray ionization quadrupole time-of-flight LC-MS, using the negative ionization mode (Agilent, Santa Clara, CA, USA).

### 2.2. SplintR qPCR assay

The probes and primers used in this study have been mentioned below:

Probe A: 5′ CTCGACCTCTCTATGGGCAGTCACGACAGGAGTCGCGCG 3′

Probe B: 5′ pCTAGGGGCCGCTGAGTCGGAGACACGCAGGGCTTAA 3′

Forward primer: 5′ GCTCGACCTCTCTATGGGC 3′

Reverse primer: 5′ TTAAGCCCTGCGTGTCTCC 3′

Double-quenched probe: 5′ /FAM/CTAGCGCGC/ZEN/GACTCCGTCGTG/IABkFQ/ 3′

Probe B has a 5′ phosphate(p) required for enzymatic ligation to Probe A. The fluorescent dye used in this study was 6-carboxyfluorescein (FAM). ZEN™was used as an internal quencher, while IABkFQ (Iowa Black FQ) was the 3′ quencher. All qPCR probes and primers used in the study were procured from IDT. The hybridization mixture was prepared as follows: 2 µl of the standard (1 µM to 1 pM) or the sample, 2 µl of 10X SplintR^®^ ligase Reaction Buffer (NEB), 2 µl of probe A (100 nM), 2 ul of probe B (100 nM), and nuclease-free water (ThermoFisher Scientific, Waltham, MA, USA) up to 12 µl. In the case of standard for biological samples, 2 µl of serum or tissue lysate was added to the hybridization mixture to form the same matrix condition. The hybridization mixture was heated to 95°C for 5 min, and then cooled to 37°C at a rate of 0.1°C per sec. After the hybridization of the probes to the target oligonucleotides, 8 µl of diluted SplintR^®^ ligase (2.5 units per reaction) was added to each hybridization mixture. Ligation of the probes was achieved by incubation at 37°C for 30 min. The ligase mixture was heat-inactivated at 65°C for 20 min.

The qPCR reaction for quantification of the ligated probes in the test article was prepared as follows: 2 µl of the ligation mixture described above, 10 µl of iTaq Super Mix (BioRad, Hercules, CA, USA), 1 µl of forward and reverse primers (10 µM), 0.5 µl double-quenched probe (10 µM), and nuclease-free water added to make up a final volume of 20 µl. The qPCR reactions were conducted in triplicate using the CFX96 Real-time System (BioRad) under the cycling conditions: initial denaturation at 95°C for 3 min, followed by 40 cycles of denaturation at 95°C for 10 sec and extension at 60°C for 30 sec.

### 2.3. 2′-O-MOE gapmer administration in mice

The administration of 2′-*O*-MOE gapmer was performed in female FVB mice (Charles River Laboratories, Wilmington, MA), 8 weeks of age, adhering to the Institutional Animal Care and Use Committee (IACUC) protocols of the University of Massachusetts Medical School. The 2′-*O*-MOE gapmer was diluted in 100 µl Phosphate Buffered Saline (PBS, ThermoFisher) (200 µM), or 100 µl PBS for control group, and was administered by tail vein injection in 5 mice per group (treatment *vs*. control). After 1 h of the initial inoculation, mice were sacrificed. The blood and tissues of the euthanized animals were harvested.

### 2.4. Preparation of serum and tissue samples

Blood was collected through cardiac puncture and allowed to stand for 1 h at room temperature for clotting without the addition of a clot activator. The blood sample was then centrifuged at 10,000xg for 5 min at room temperature. The serum was collected from the supernatant.

A sample of 20-30 mg of different tissues (liver, kidney, lung, heart, muscle, and brain) were removed from the mice treated with PBS or 2′-*O*-MOE gapmer, respectively and weighed followed by tissue disruption using Tissuelyser II (Qiagen, Waltham, MA, USA) in 30 µl mg^-1^ RIPA buffer (Alfa Aesar, Haverhill, MA, USA) containing 10 mM Tris-HCl (pH 7.4), 150 mM NaCl, 1% NP-40, 0.5% sodium deoxycholate, and 0.1% sodium dodecyl sulfate with 5 mM EDTA and 1 mM EGTA. Tissues were incubated for 1h on ice with vortexing every 10 min, followed by centrifugation at 16,000xg for 30 min at 4°C. The supernatant was collected into a new 1.5-ml microcentrifuge tube and stored at -80°C until subsequent analysis for the presence of 2′-*O*-MOE gapmer was performed.

### 2.5. Data analysis

Mean, standard error mean, linear regression, and statistical tests were performed using GraphPad Prism V 8.4.2 (GraphPad Software, San Diego, CA, USA). The regression line was calculated using the log of standard concentration in the sample *vs*. the mean quantification cycle (Cq). The P-value in tissue quantification was calculated using the Two-stage linear step-up procedure of Benjamini, Krieger, and Yekutieli with Q=1%.

## Results and Discussion

Reliable and relatively straightforward bioanalytical assays for the detection and quantification of therapeutic ASOs will help in understanding the mechanism of action of oligonucleotides, which will facilitate their translation from preclinical to clinical settings. This study was aimed at developing a quick and reliable new method to detect and quantify ASOs in biological samples. Since enzymatic ligation with the *Chlorella* virus DNA ligase (SplintR^®^ ligase) [24,25] is a quick and cost-effective method to detect miRNAs, the objective of this study was to optimize and apply this enzymatic probe ligation-based qPCR technique to detect and quantify ASOs.

The quantification of ASOs was achieved using the SplintR qPCR assay (**Fig. 1A**). Each probe consisted of 3 parts: oligonucleotide-binding (OB), linker, and primer-binding (PB) regions. For hybridization, the oligonucleotide-binding region had a sequence complementary to the target ASO. The oligo-binding region was separated from the primer-binding site by the linker region of the probe, thus extending the length of the ligation product for efficient PCR amplification. The primer-binding regions were designed to be complementary to qPCR primers. The oligonucleotide-binding region varied depending on the target ASO, while the linker and primer-binding regions remained constant. Probe A was synthesized with a 5′-phosphoryl group, while probe B had a 3′-hydroxyl group, which is essential for successful ligation. After probe A and B were hybridized to the target ASO, SplintR^®^ ligase was used to ligate the nick between probe A and B. As the SplintR^®^ ligase had a higher activity to the probe that was hybridized to the target ASO (compared with unhybridized probes), the ligated product increased proportionally with the amount of ASOs available in the sample (39). For specific and sensitive qPCR analysis, we designed a double-quenched qPCR probe, which relied on exonucleolytic release of a 5′ fluorophore **(Fig.1B)**.

**Figure 1.**
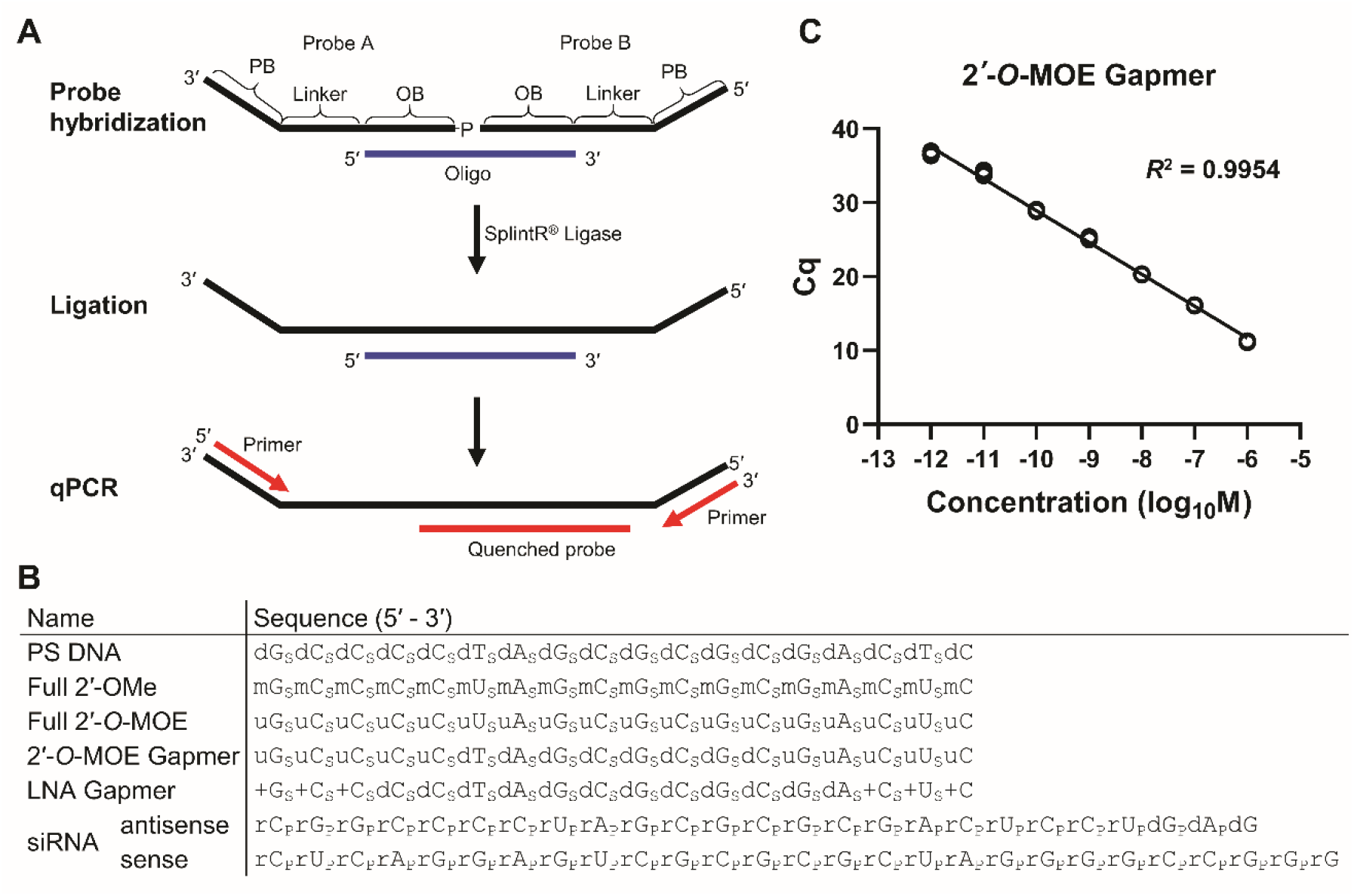
**(A)** Schematic diagram of antisense oligonucleotide (ASO) quantification using the SplintR qPCR assay. Probe A and B are hybridized to the target oligonucleotide by complementary oligonucleotide-binding (OB) sequence. After hybridization, two probes are ligated by SplintR^®^ ligase followed by qPCR with primers that recognize the primer-binding (PB) sequence, and detect the amplification using the quenched probe. **(B)** Summary of ASOs used in this study. Oligonucleotide modifications have been represented as subscripts: p - phosphodiester linkage; s - phosphorothioate linkage; dN - DNA; mN - 2′-OMe RNA; uN - 2′-*O*-MOE RNA; +N - LNA; rN - RNA. **(C)** Quantification of 2′-*O*-MOE gapmer. Standard curve plotted from Cq value *vs*. the log of 2′-*O*-MOE gapmer concentration in the sample.

To determine whether the SplintR qPCR assay can be used to quantify modified ASOs, the 2′-*O*-MOE gapmer with a PS linkage was tested thereafter. We quantified a dilution series of 2′-*O*-MOE gapmer in water to generate the standard curves (**Supplementary Fig. S1**). The SplintR qPCR assay showed a broad range of linearity over 7 orders of magnitude from 1 pM to 1 µM, with the coefficient of determination *R*^2^ > 0.99. The 2′-*O*-MOE gapmer ASO was detected at concentration as low as 1 pM, demonstrating high sensitivity of the quantitative method described in this study (**Fig.1C**).

Preliminary tests to evaluate the efficacy of SplintR qPCR assay to quantify ASOs in biological samples involved preparation of a dilution series of 2′-*O*-MOE gapmer in mouse serum and mouse liver lysate, respectively, which was followed by a subsequent round of the SplintR qPCR assay. Our results show that SplintR qPCR assay allowed efficient and sensitive detection of 2′-*O*-MOE gapmer at low concentrations ranging from 1 pM to 1 µM in serum and liver lysate-based samples, with the coefficient of determination *R*^2^ > 0.99 (**Supplementary Fig. S2**).

In the subsequent experiments the efficiency of SplintR qPCR assay in quantifying the ASOs administered to mice via tail vein injection was determined. The test animals were administered with 2′-*O*-MOE gapmer (20 nmol) by tail vein injection. After 1 h the animals were euthanized and serum, liver, kidney, lung, heart, muscle, and brain samples from each mouse were collected for subsequent quantification of 2′-*O*-MOE gapmer using the SplintR qPCR assay (**Fig. 2**). Negative controls included samples from mice injected with PBS alone. We generated standard curves for 2′-*O*-MOE gapmer diluted in serum or tissue lysates: serum, liver, and kidney with linear detection across 7 orders of magnitude ranging from 1 pM to 1 µM; lung, heart, and muscle, with linear detection across 6 orders of magnitude ranging from 10 pM to 1 µM (**Supplementary Fig. S3**). The SplintR qPCR assay could only detect the 2′-*O*-MOE gapmer in brain lysate across 5 orders of magnitude from 100 pM to 1 µM (**Supplementary Fig. S3**). The assay has reduced sensitivity for brain lysate samples, which may be attributed to the high lipid content of the brain tissue. Nevertheless, the results presented in this study indicate that the SplintR qPCR assay is compatible with biological samples and efficiently detects ASOs. The SplintR qPCR assay revealed that the 2′-*O*-MOE gapmer levels were abundant in serum (437.3 pmol ml^-1^), liver (1.2 pmol mg^-1^), and kidney (12.7 pmol mg^-1^), respectively. Lower levels were detected in lung (386.0 fmol mg^-1^), heart (305.3 fmol mg^-1^), and muscle (411.5 fmol mg^-1^), respectively. The quantification assay in brain tissue showed no detection above of background levels which may be attributed to lower transfer across the blood-brain barrier.

**Figure 2.**
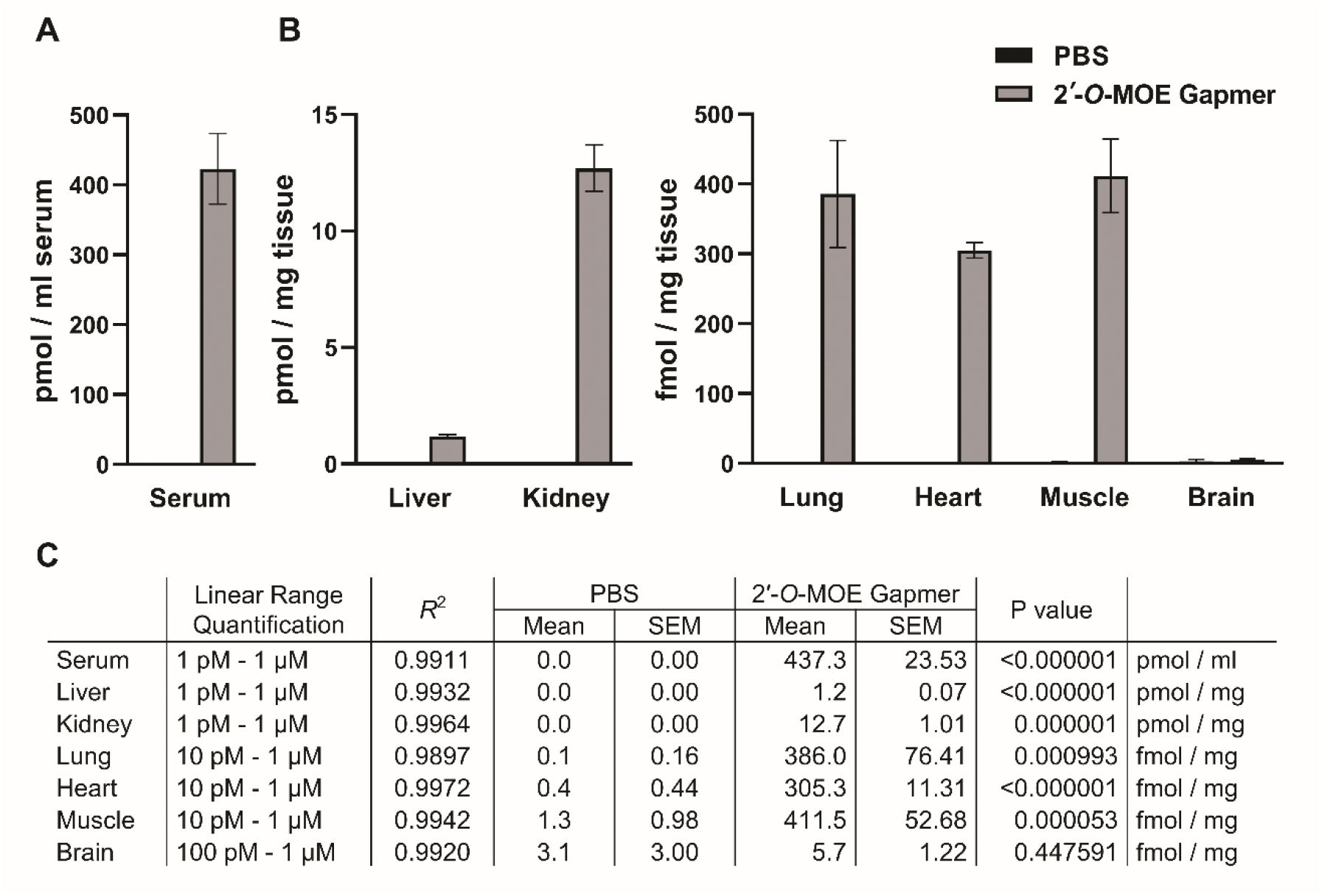
Detection of 2′-*O*-MOE gapmer after tail vein injection in a mouse model. Blood and tissues were collected 1 h after administration of 20 nmol 2′-*O*-MOE gapmer or PBS (control). Serum was purified from blood, and tissues were lysed using RIPA buffer. The concentration of 2′-*O*-MOE gapmer in serum **(A)** and tissues **(B)** was quantified using the SplintR qPCR assay. **(C)** The table lists the concentration of the 2′-*O*-MOE gapmer and the linear range of the standard curve. n=5 mice per group, Mean±Standard Error Mean (SEM).

In order to understand the scope of applications of the SplintR qPCR assay discussed in this study, we tested other types of ASOs. This technique was tested for various types of ASOs with chemical modifications including uniformly modified ASOs (PS DNA, 2′-OMe, 2′-*O*-MOE), gapmers (2′-*O*-MOE, LNA), and siRNA, respectively. Our results yieldeddetection even for highly modified full 2′-*O*-MOE although this substrate required a higher amount of SplintR ligase in the ligation step.

Quantification assays performed for fully PS-modified DNA yielded positive detection outcomes across a linear range of 7 orders of magnitude from 1 pM to 1 µM. In contrast, the fully modified 2′-OMe and 2′-*O*-MOE ASOs were detected over 6 orders of magnitude from 100 pM to 10 µM (**Fig. 3A**). In order to obtain a linear range of detection of the fully modified 2′-*O*-MOE ASO 10-fold higher amounts of ligase (25 units/reaction) were required (**Supplementary Fig. S4**). The bulky modification of 2’-O-MOE RNA seemed to partially inhibit the enzymatic ligation reaction, requiring some optimization of both ligation probe sequences and reaction conditions.

**Figure 3.**
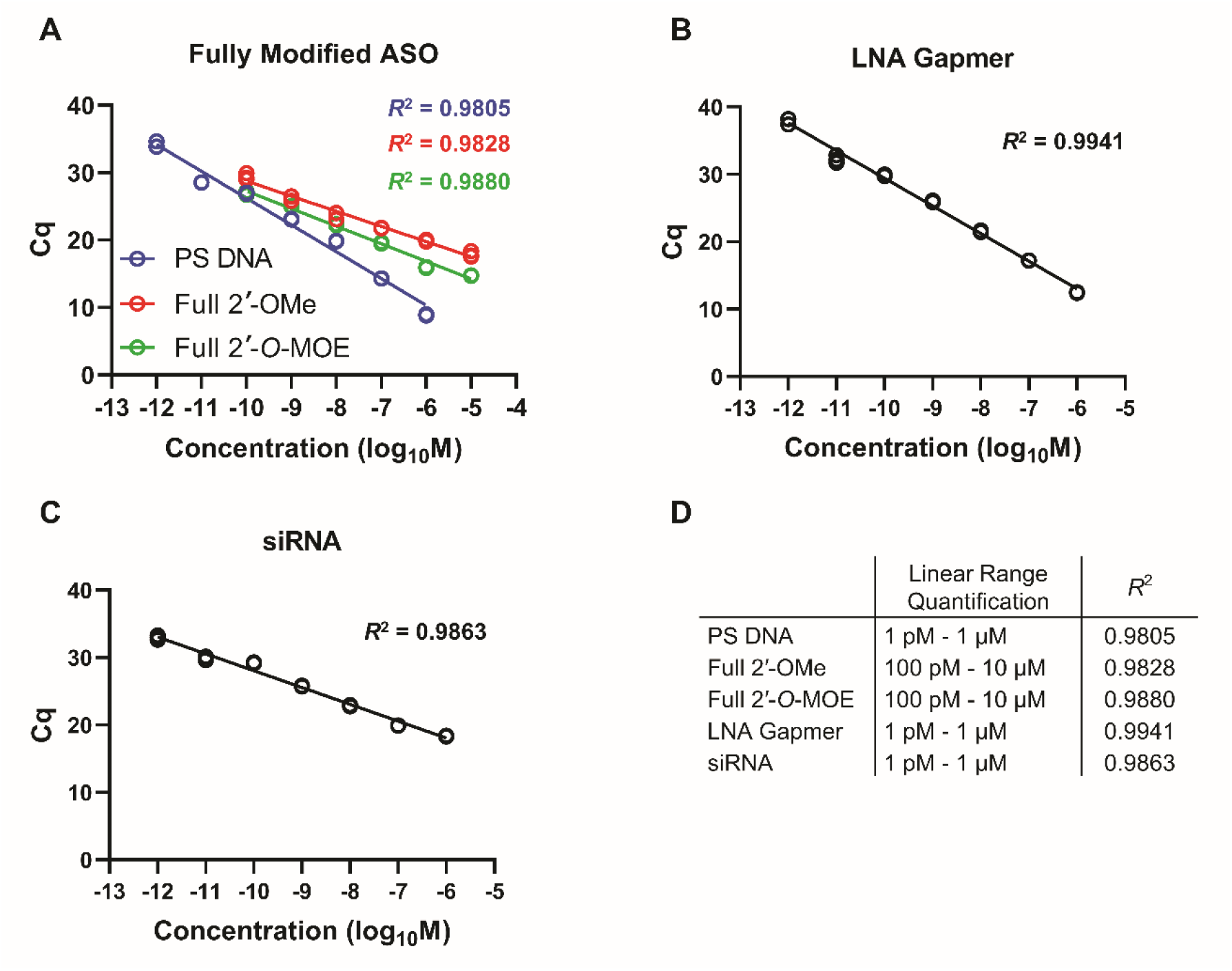
Quantification of ASOs using the SplintR qPCR assay. Various ASOs were detected, which includes: **(A)** fully modified ASOs, **(B)** LNA gapmer, **(C)** siRNA, and **(D)** enlists the parameters.

Next, we tested a LNA gapmer widely used for RNase H mediated gene silencing along with a 2′-*O*-MOE gapmer. The LNA gapmer showed optimum quantification over 7 orders of magnitude from 1 pM to 1 µM (**Fig. 3B**), similar to that obtained for 2′-*O*-MOE gapmer. We also tested siRNA, which resulted in the linear range quantification over 7 orders of magnitude from 1 pM to 1 µM (**Fig. 3C**). The specifics of the linear range quantification and the coefficient of determination (*R*^2^) has been presented in **Fig. 3D**.

To improve platform technologies for oligonucleotide therapeutics and design apply them to new diseases, quick and cost-effective techniques to detect and quantify therapeutic ASOs are pertinent. The SplintR qPCR assay does not require specialized equipment, but uses qPCR reagents and equipment available to most molecular biology laboratories. The SplintR qPCR assay has several advantages over the previously described chemical ligation qPCR strategy[23]. The *Chlorella* virus SplintR^®^ ligase is readily available and can be used directly without any special modifications of the splint ligation probes, and the procedure is faster. Thus, the SplintR qPCR assay is a simple and cost-effective way to detect and quantify therapeutic ASOs in biological samples, such as serum or biopsy-based tissues. The dynamic range of the assay is superior of that of hybridization ELISA, and will be easier to implement for many laboratories. Therefore, the SplintR qPCR assay described in this study is likely to become a useful tool for groups studying therapeutic development and pharmacokinetics of oligonucleotide therapeutics.

## Acknowledgments

The authors thank Dr. Darryl Conte Jr. for careful review of the article and for helpful discussions.

## Author Disclosure Statement

J.K.W. is Scientific Advisory Board member of PepGen and ad hoc consultant for BridgeBio and Flagship Pioneering.

## Funding Information

This work was supported by the Ono Pharma Foundation (Breakthrough Science Award to J.K.W.) and the National Institute of Neurological Disorders and Stroke, the National Institutes of Health (R01-NS111990 to J.K.W.).

## Figure Legends

**Supplementary Figure S1.**
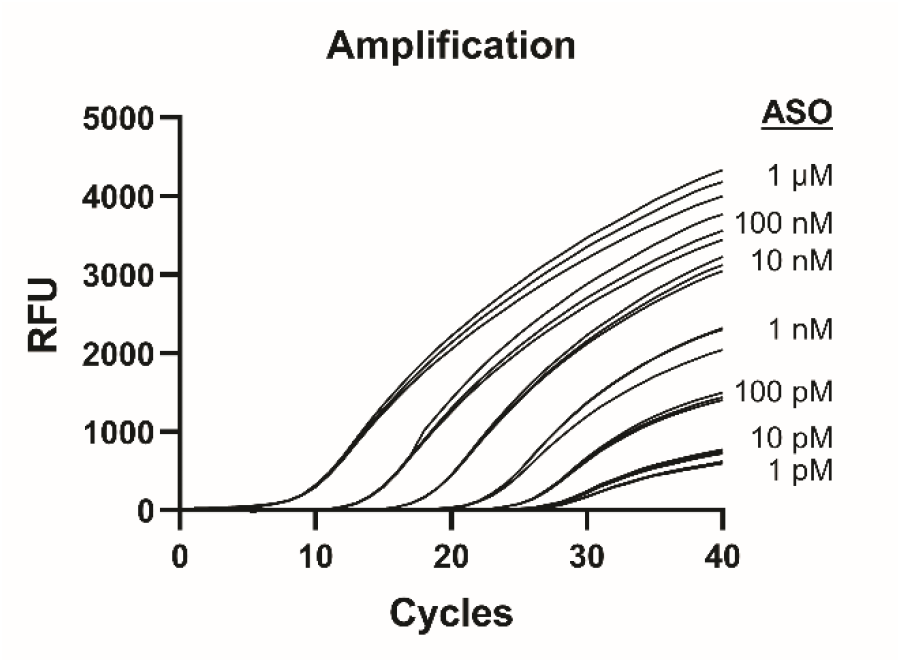
Quantification of 2′-*O*-MOE gapmer. Amplification curve of the SplintR qPCR assay of a 2′-*O*-MOE gapmer serially diluted in water from 1 pM to 1 µM. The concentration of the 2′-*O*-MOE gapmer is written at the right of the amplification curve.

**Supplementary Figure S2.**
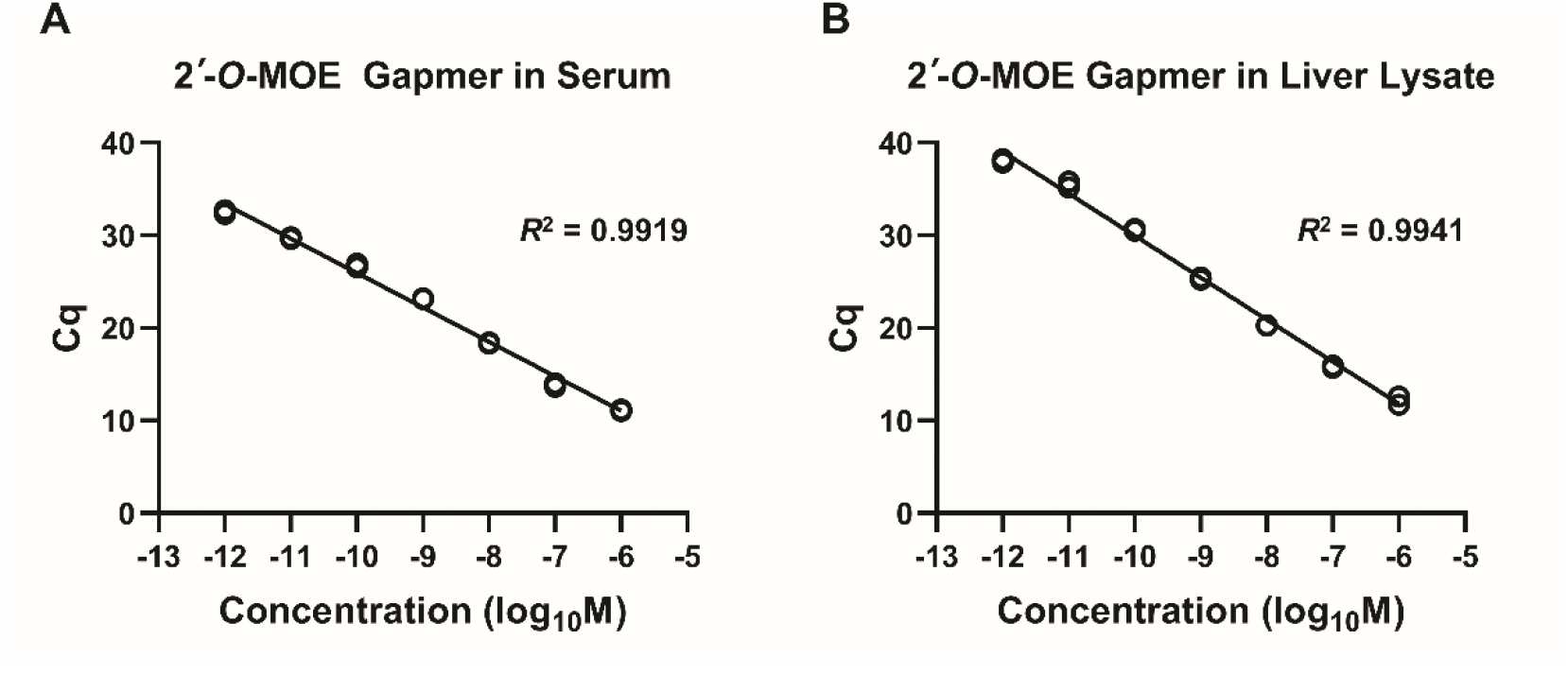
Quantification of 2′-*O*-MOE gapmer in biological samples. The standard curve was generated using diluted 2′-*O*-MOE gapmer fortified with mouse **(A)** serum or **(B)** liver lysate.

**Supplementary Figure S3.**
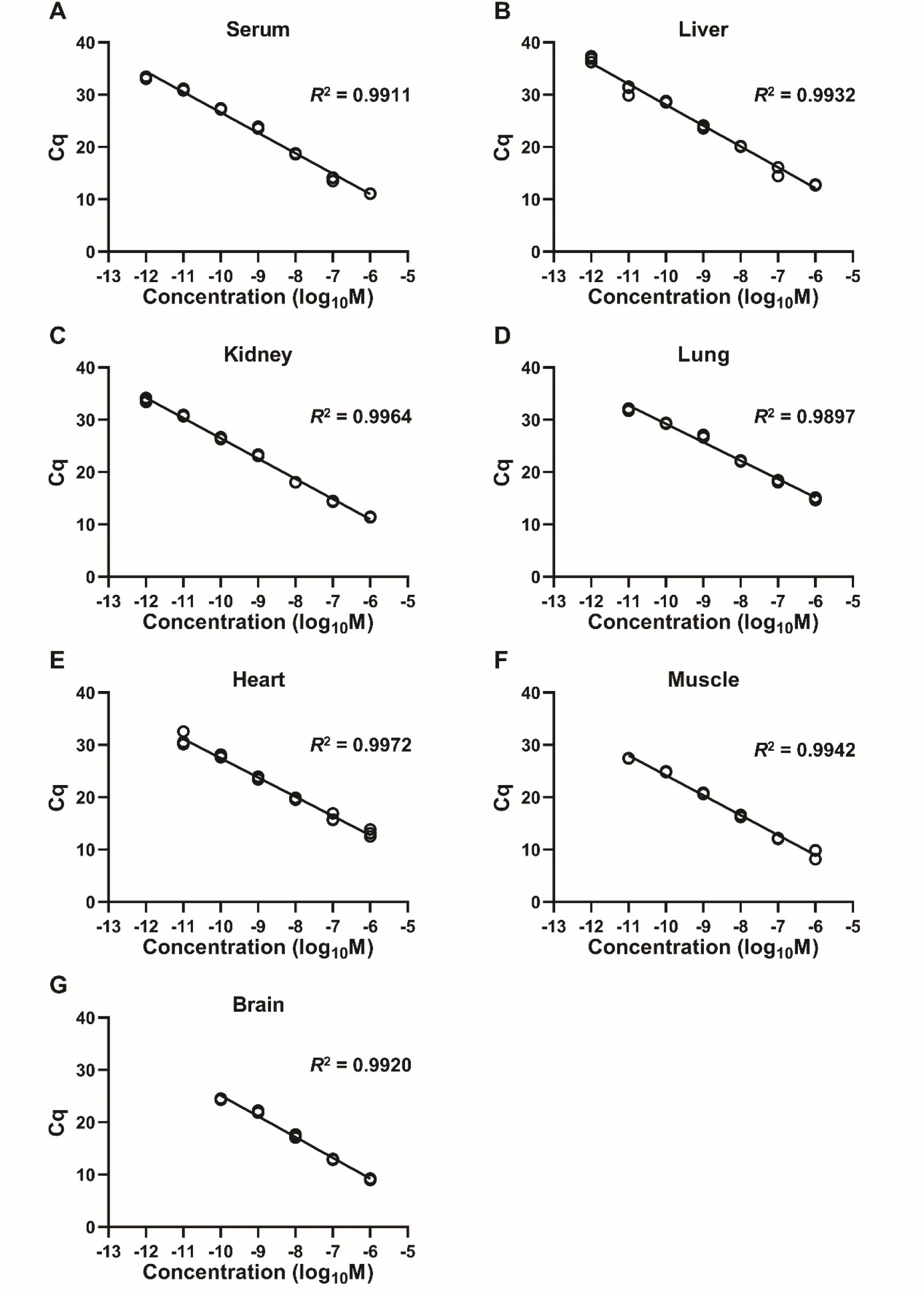
Standard curve for 2′-*O*-MOE gapmer from: **(A)** serum, and **(B-G)** different tissue lysates.

**Supplementary Figure S4.**
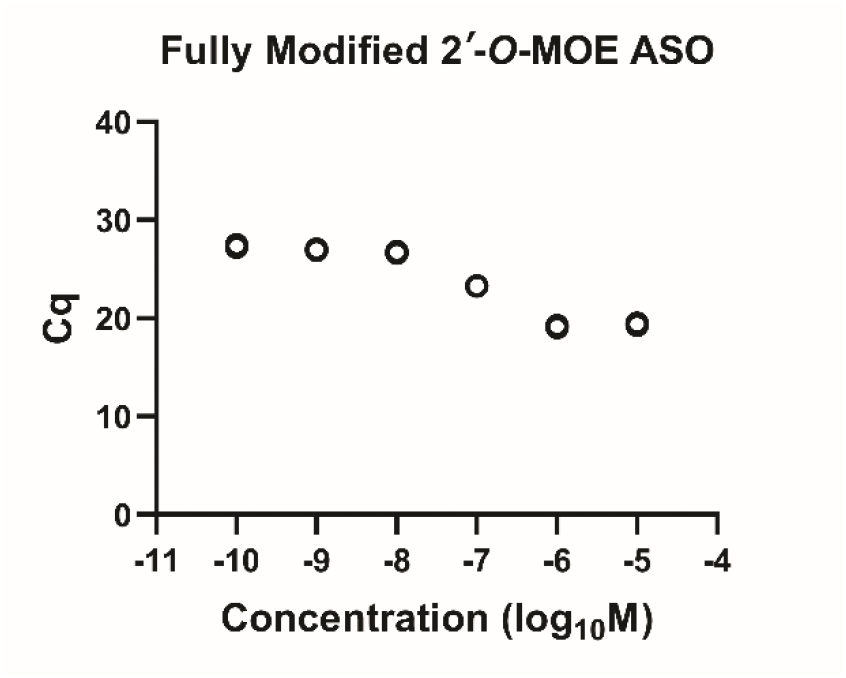
Standard curve for fully modified 2′-*O*-MOE oligonucleotide using 2.5 units/reaction of SplintR^®^ ligase.

## References

1. Stephenson ML and PC Zamecnik. (1978). Inhibition of Rous sarcoma viral RNA translation by a specific oligodeoxyribonucleotide. Proc Natl Acad Sci U S A 75:285–288.

2. Zamecnik PC and ML Stephenson. (1978). Inhibition of Rous sarcoma virus replication and cell transformation by a specific oligodeoxynucleotide. Proc Natl Acad Sci U S A 75:280–284.

3. Yamamoto T, M Nakatani, K Narukawa and S Obika. (2011). Antisense drug discovery and development. Future Med Chem 3:339–365.

4. Khvorova A and JK Watts. (2017). The chemical evolution of oligonucleotide therapies of clinical utility. Nat Biotechnol 35:238–248.

5. Sharma VK and JK Watts. (2015). Oligonucleotide therapeutics: chemistry, delivery and clinical progress. Future Med Chem 7:2221–2242.

6. Saleh AF, AA Arzumanov and MJ Gait. (2012). Overview of alternative oligonucleotide chemistries for exon skipping. Methods Mol Biol 867:365–378.

7. Eckstein F. (2014). Phosphorothioates, essential components of therapeutic oligonucleotides. Nucleic Acid Ther 24:374–387.

8. Crooke ST, TA Vickers and XH Liang. (2020). Phosphorothioate modified oligonucleotide-protein interactions. Nucleic Acids Res 48:5235–5253.

9. Monia BP, EA Lesnik, C Gonzalez, WF Lima, D McGee, CJ Guinosso, AM Kawasaki, PD Cook and SM Freier. (1993). Evaluation of 2’-modified oligonucleotides containing 2’-deoxy gaps as antisense inhibitors of gene expression. J Biol Chem 268:14514–14522.

10. Prakash TP and B Bhat. (2007). 2’-Modified oligonucleotides for antisense therapeutics. Curr Top Med Chem 7:641–649.

11. Greenberger LM, ID Horak, D Filpula, P Sapra, M Westergaard, HF Frydenlund, C Albæk, H Schrøder and H Ørum. (2008). A RNA antagonist of hypoxia-inducible factor-1α, EZN-2968, inhibits tumor cell growth. Mol Cancer Ther 7:3598–3608.

12. Bonham MA, S Brown, AL Boyd, PH Brown, D. Bruckenstein, JC Hanvey, SA Thomson, A Pipe, F Hassman, JE Bisi and et al. (1995). An assessment of the antisense properties of RNase H-competent and steric-blocking oligomers. Nucleic Acids Res 23:1197–1203.

13. Aartsma-Rus A, L van Vliet, M Hirschi, AA Janson, H Heemskerk, CL de Winter, S de Kimpe, JC van Deutekom, PA t Hoen and GJ van Ommen. (2009). Guidelines for antisense oligonucleotide design and insight into splice-modulating mechanisms. Mol Ther 17:548–553.

14. Monia BP, EA Lesnik, C Gonzalez, WF Lima, D McGee, CJ Guinosso, AM Kawasaki, PD Cook and SM Freier. (1993). Evaluation of 2’-modified oligonucleotides containing 2’-deoxy gaps as antisense inhibitors of gene expression. J Biol Chem 268:14514–14522.

15. Rinaldi C and MJA Wood. (2018). Antisense oligonucleotides: the next frontier for treatment of neurological disorders. Nat Rev Neurol 14:9–21.

16. Wang F, T Zuroske and JK Watts. (2020). RNA therapeutics on the rise. Nat Rev Drug Discov.

17. Deprey K, N Batistatou and JA Kritzer. (2020). A critical analysis of methods used to investigate the cellular uptake and subcellular localization of RNA therapeutics. Nucleic Acids Res 48:7623–7639.

18. Basiri B and MG Bartlett. (2014). LC-MS of oligonucleotides: applications in biomedical research. Bioanalysis 6:1525–1542.

19. Kaczmarkiewicz A, Ł Nuckowski, S Studzińska and B Buszewski. (2019). Analysis of Antisense Oligonucleotides and Their Metabolites with the Use of Ion Pair Reversed-Phase Liquid Chromatography Coupled with Mass Spectrometry. Crit Rev Anal Chem 49:256–270.

20. Verhaart IE, CL Tanganyika-de Winter, TG Karnaoukh, IG Kolfschoten, SJ de Kimpe, JC van Deutekom and A Aartsma-Rus. (2013). Dose-dependent pharmacokinetic profiles of 2’-O-methyl phosphorothioate antisense oligonucleotidesin mdx mice. Nucleic Acid Ther 23:228–237.

21. Thayer MB, SC Humphreys, KS Chung, JM Lade, KD Cook and BM Rock. (2020). POE Immunoassay: Plate-based oligonucleotide electro-chemiluminescent immunoassay for the quantification of nucleic acids in biological matrices. Sci Rep 10:10425.

22. Wong ML and JF Medrano. (2005). Real-time PCR for mRNA quantitation. Biotechniques 39:75–85.

23. Boos JA, DW Kirk, ML Piccolotto, W Zuercher, S Gfeller, P Neuner, A Dattler, WL Wishart, F Von Arx et al. (2013). Whole-body scanning PCR; a highly sensitive method to study the biodistribution of mRNAs, noncoding RNAs and therapeutic oligonucleotides. Nucleic Acids Res 41:e145.

24. Jin J, S Vaud, AM Zhelkovsky, J Posfai and LA McReynolds. (2016). Sensitive and specific miRNA detection method using SplintR Ligase. Nucleic Acids Res 44:e116.

25. Wee EJH and M Trau. (2016). Simple Isothermal Strategy for Multiplexed, Rapid, Sensitive, and Accurate miRNA Detection. ACS Sens 1:670–675.

26. Pendergraff HM, PM Krishnamurthy, AJ Debacker, MP Moazami, VK Sharma, L Niitsoo, Y Yu, YN Tan, HM Haitchi and JK Watts. (2017). Locked Nucleic Acid Gapmers and Conjugates Potently Silence ADAM33, an Asthma-Associated Metalloprotease with Nuclear-Localized mRNA. Mol Ther Nucleic Acids 8:158–168.

